# Sex differences in transcription-associated mutagenesis in the human germline

**DOI:** 10.1101/2025.09.18.677082

**Authors:** Minyoung J. Wyman, Ipsita Agarwal, Marc de Manuel, Natanael Spisak, Molly Przeworski

## Abstract

In humans, germline mutation rates are three- to four-fold higher in males than females, for largely unknown reasons. We investigated whether transcription, a well-documented source of both DNA damage and repair in somatic tissues, is associated with sex differences in germline mutations. To this end, we used expression data from male and female germline cells and phased *de novo* germline mutations from pedigrees. Focusing on protein-coding genes, we found no relationship between the male mutation rate and gene expression levels in the fetal germline or in adult testis tissue, despite evidence for transcriptional asymmetry. Individual stages of spermatogenesis differ in their contribution to mutation, however: expression levels in spermatogonial stem cells are significantly positively associated with paternal mutation rates, while those in primary spermatocytes are significantly negatively associated. Thus, transcription may have varying effects over male gametogenesis that are not readily detected from its cumulative effect on the total germline mutation rate. In females, by contrast, mutation rates increase significantly with transcription levels in the fetal germline, adult oocytes and adult ovary tissues, consistent with widespread transcription asymmetry. We confirm the difference between the sexes by analyzing phased mutations from three-generation pedigrees and the lack of an association in males by analyzing paternal mutations from seminiferous tubules and sperm. Thus, transcription has distinct effects on the mutation rate in the two sexes, leading to an increase in mutations in females but not males, in contrast to what one might expect from the overall paternal bias in germline mutations.

## Introduction

Mutations in the human male and female germlines differ in a number of regards. Notably, the rate of *de novo* mutation (DNM) is three to four times higher in males than in females (Jónsson et al. 2017), and the mutational spectrum differs measurably between the sexes (Goldmann et al. 2016; Jónsson et al. 2017; Agarwal and Przeworski 2019; Spisak et al. 2024). The basis for these sex-specific differences remains poorly understood, but multiple lines of evidence suggest that germline point mutations in both sexes arise predominantly from DNA damage that is inaccurately repaired (Gao et al. 2016; Gao et al. 2019; Seplyarskiy et al. 2019; Seplyarskiy et al. 2021; de Manuel et al. 2022; Spisak et al. 2024). Therefore, the sex difference in mutations may reflect, at least in part, differences in damage rates or repair accuracies.

Transcription is a known source of both DNA damage and repair, shaping mutation rates in human somatic tissues (Haradhvala et al. 2016; Pleasance et al. 2010; Alexandrov et al. 2013; Lodato et al. 2015; Alexandrov et al. 2020; Jeffries et al. 2025). High transcriptional levels can create supercoiled DNA in front of or behind the advancing RNA polymerase II (RNAPII) (Wang 2002). TOP1 relieves positive supercoiling by nicking the DNA backbone, but cleavage at sites of genome-embedded ribonucleotides can lead to short deletions (Takahashi et al. 2011), a signature detected in the human germline (Reijns et al. 2022). Negative supercoiling results in single stranded DNA, a chemically less stable conformation (Jinks-Robertson and Bhagwat 2014), which introduces opportunities for lesions on both strands (Gaillard and Aguilera 2016; Cho and Jinks-Robertson 2017).

On the other hand, transcription can recruit the repair machinery. Transcription-coupled repair (TCR), a sub-pathway of nucleotide excision repair, may intervene when a bulky lesion on the transcribed strand stalls RNAPII (Hanawalt and Spivak 2008; Vermeulen and Fousteri 2013; Spivak 2015; Nicholson et al. 2024). TCR also processes R-loops (Sollier et al. 2014; Sollier and Cimprich 2015). In the soma, TCR repairs insertions and deletions on the transcribed strand in conjunction with mismatch repair (Georgakopoulos-Soares et al. 2020). TCR cross-talks with the components of base excision repair (BER) to repair smaller oxidative lesions encountered during transcription (Svejstrup 2002; Kim and Jinks-Robertson 2010; Chakraborty et al. 2021). In addition, a separate process known as domain-associated repair can repair either strand within transcribed regions (Nouspikel et al. 2006; Zheng et al. 2014). The effect of transcription levels on the overall mutation rate (i.e., across both strands) will therefore depend upon the combination of all repair and damage-inducing processes.

In most somatic cell types, the mutation rate appears to be negatively correlated with gene expression levels, which suggests that the effect of TCR dominates over transcription-related damage (Pleasance et al. 2010; Chapman et al. 2011; Chen et al. 2017); among post-mitotic neurons, however, the mutation rate appears positively correlated to expression (Lodato et al. 2015; Jeffries et al. 2025). The patterns in the germline are less clear: previous studies have suggested a negative (Xia et al. 2020; Xia and Yanai 2022), a positive (Chen et al. 2017; Liu and Zhang 2020), or no significant correlation (Moore et al. 2021) between mutation rates and expression levels in the male germline. The discrepant findings among studies may stem in part from differences in the male expression data used: e.g., bulk adult testis tissue (Chen et al. 2017), seminiferous tubules (Moore et al. 2021), or single-cell adult spermatogonia (Xia et al. 2020; Liu and Zhang 2020). Studies also differed in whether they considered only mutations in testes (Moore et al. 2021), all germline mutations (Xia et al. 2020; Liu and Zhang 2020), or paternally and maternally phased mutations separately (Chen et al. 2017), as well as in what covariates were included, and whether genes with no mutations were included in the analyses (Xia et al. 2020; Xia and Yanai 2022; Liu and Zhang 2020). Finally, to date, only one study looked specifically at maternal germline mutations, reporting a positive correlation with adult bulk ovary tissue expression (Chen et al. 2017).

Male and female germlines are a succession of distinct cell types (Figure 1) with dynamic transcriptional programs that differ between the sexes. For instance, in the fetus, female germ cells have higher levels of transcription associated with meiosis compared to similarly staged male germ cells (Garcia-Alonso et al. 2022). Moreover, within the germlines of each sex, expression is globally lower in some cell types (e.g., primordial oocytes and spermatogonial stem cells) compared to others (e.g., antral oocytes and primary spermatocytes) (Guo et al. 2018; Zhang et al. 2018). Such broad-scale differences suggest that the effect of transcription on mutation rates could plausibly vary between sexes, as well as across development within a sex, depending on the particular balance of repair and damage associated with a cell type.

**Figure 1.**
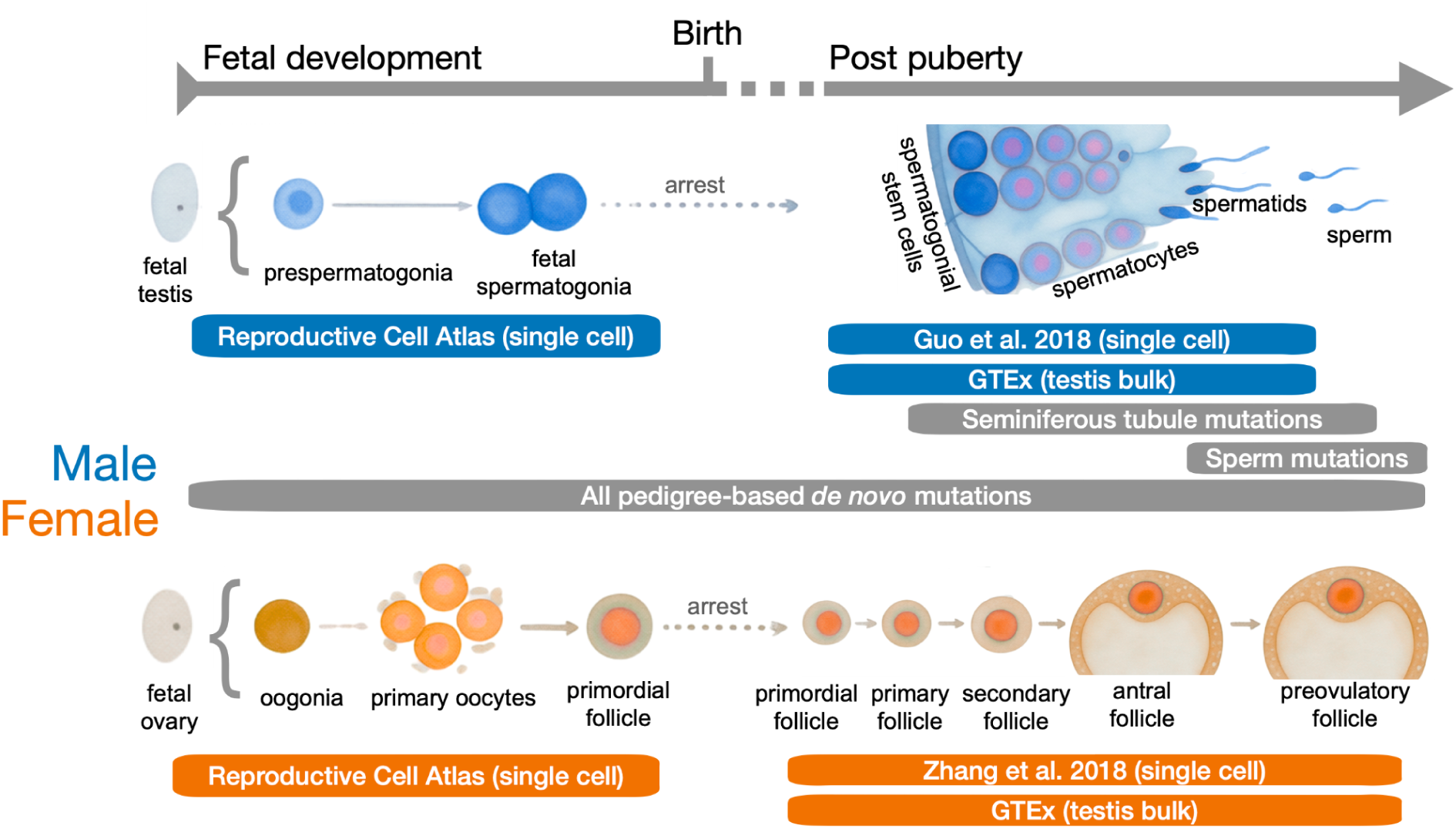
Data sources with respect to male and female germline development across fetal and adult stages. The span of the text bars indicate the developmental stages from which the four gene expression datasets and three mutation datasets used in this study were derived. This figure was adapted from (Saitou et al. 2026).

Although the relationship between germline expression levels and mutation rates remains unclear, indirect evidence points to an effect of transcription on human germline mutations. Specifically, strand asymmetry in transcribed regions (i.e., T-asymmetry) indicates differential repair or damage rates between the DNA strands for at least some mutation types: there are fewer A>G substitutions on the transcribed strand than on the non-transcribed strand in mammalian comparisons (Green et al. 2003), and in analyses of DNM and single nucleotide variant data from current human populations (Xia et al. 2020; Chen et al. 2017; Seplyarskiy et al. 2021; Seplyarskiy and Sunyaev 2021). Thus, expression appears to have effects on either damage or repair rates in the germline.

With these considerations in mind, we set out to clarify how transcription-related repair and damage affect human germline mutations, using DNMs identified in pedigrees, seminiferous tubules, and sperm, as well as using both fetal and adult expression data (Figure 1). Specifically, we characterized the relationship between mutation rates and protein-coding expression levels in the germline of both sexes.

## Materials and methods

### Mutation rates in transcribed regions

We combined 451,005 single nucleotide DNMs from three WGS studies that analyzed 3,792 trios (816 trios from (Goldmann et al. 2016) and 2,976 trios from (Halldorsson et al. 2019)) and 1,902 quartets (An et al. 2018), respectively. Among these mutations, 145,999 were assigned to either the paternal (115,093) or maternal (30,906) genome through read-based phasing or third generation transmission. We removed 896 mutations with reference alleles that differed between GRCh37 and GRCh38.

We also compiled 23,470 DNMs identified by sequencing seminiferous tubules from 14 donors (Moore et al. 2021), as well as 10,697 DNMs detected from sperm sequencing of 88 donors (Abascal et al. 2021; Neville et al. 2025; Shoag et al. 2025). From this set of male germline mutations, we removed five mutations with reference alleles differing between GRCh37 and GRCh38.

Among the set of mutations described above, we retained only those falling in the protein-coding genes, which are transcribed by RNAPII, the enzyme integral to initiating TCR (Hanawalt and Spivak 2008; Vermeulen and Fousteri 2013). We used only autosomal genes to avoid difficulties in interpretation stemming from changes in expression levels during male meiotic sex chromosome inactivation and female X chromosome reactivation. To avoid biasing our estimates, we included genes with no mutations, as long as they were present in the intersection of the four gene expression datasets described below (GTEx Consortium 2013; Guo et al. 2018; Zhang et al. 2018; Garcia-Alonso et al. 2022) and had a replication timing value (Ding et al. 2021) (see *Additional covariates* section), resulting in 15,220 genes total. We selected genes designated as “protein-coding” in the GENCODE annotation (GRCh38, release 41), and then selected for “transcripts” in the feature column of the comprehensive gene annotation GTF for chromosomes, which includes exons, introns, and UTRs.

We took the coordinates of all transcripts belonging to one gene and merged them to construct a single contiguous transcribed region per gene (bedtools merge command). Within the contiguous transcript, we then retained only base pairs that had at least 10X mean coverage, as reported in gnomAD v3. We chose this threshold by comparing the effect of alternative filters. A 10X filter removed <1% of all mutations and all surveyed sites whereas a more stringent 20X filter removed ∼2% of sites and eliminated 14 genes entirely. We concluded that the 10X filter balanced the need to retain potentially informative genes while also excluding positions that may not be consistently accessible by whole genome sequencing in the DNM studies represented above. Finally, the total number of base pairs in these contiguous 10X regions provided the estimate of mutational opportunities (e.g., gene length) through a logged offset term in all of the regression models and was used as the denominator in the mutation rate calculation.

Among the mutations identified in pedigree data, 194,356 were present in the 10X filtered regions of the final set of 15,220 protein-coding genes. Of these, 47,787 were assigned to the male germline and 12,712 to the female germline based on nearby variants in reads or third generation phasing (4,865 were inferred to be paternal and 1,136 maternal using a third generation specifically). In turn, 9,869 mutations from seminiferous tubules and 4,613 from sperm fell within the filtered regions.

To explore the association of expression levels with different mutation types, we consider transitions at CpG sites and all other mutation types separately, corresponding roughly to mutational signatures SBS1 and SBS5 respectively, the two main signatures prevalent among germline mutations (Rahbari et al. 2016; Moore et al. 2021).

### Gene expression levels

Given the differences in transcription between sexes and across development, we used four sets of gene expression data, corresponding to the expression levels in fetal and adult stages (Figure 1). Two datasets had male and female expression data collected and analyzed using similar methods, enabling a direct comparison of the sexes: single-cell RNA-seq using fetal germ cells from the Reproductive Cell Atlas (Garcia-Alonso et al. 2022) and gonadal bulk tissue from adults (GTEx Consortium 2013).

Specifically, the Reproductive Cell Atlas is composed of single cell germline expression from 16 male and 28 female fetal donors (Garcia-Alonso et al. 2022). The UMI counts from various stages in males (primordial germ cells (PGCs), germ cells, prespermatogonia) and females (PGCs, germ cells, oogonia, oocytes) were pseudobulked (Squair et al. 2021; Murphy and Skene 2022) for each gene and within each donor separately to preserve the biological replication at the donor-level (Zimmerman et al. 2021). Then the donor-level, gene-specific sums were normalized by the total number of reads in the relevant donor, converted to counts per million (CPM) and incremented by 1 (Aitchison 1986), so that genes with zero expression could be log-transformed (base=2) and included. The pseudocount is essentially a weak prior that truly zero expression is unlikely in compositional data (Robinson et al. 2010; Friedman and Alm 2012; Fernandes et al. 2014; Love et al. 2014; Satija et al. 2015); moreover, a pseudocount permits the inclusion of genes that have sex-limited expression so that males and females can be compared for those genes. The donor-specific log_2_(CPM+1) values were then averaged within each sex to obtain one measure of expression per gene. This pseudobulking procedure is meant to produce a measure of fetal germline expression similar in nature to bulk tissue expression obtained from traditional RNAseq. The normalized expression values on the same scale enable us to test for sex differences at the fetal stage.

In turn, the GTEx expression data came from 361 bulk testis and 180 bulk ovary tissue samples. Bulk gonadal expression data, being comprised of both somatic and germline cells, provide a noisy estimate of transcription levels from adult gametes. Nevertheless, if germline transcription is a main source of variation in germline mutation rates for protein-coding genes, we do not expect expression levels from somatic tissues to distort the germline effects (see also *Stage-specific gene expression* section). The raw read counts for the ovary and testis samples were jointly TMM-normalized using edgeR (with default setting prior.count=0.25) and converted to CPM. The log_2_CPM values were averaged across all samples within each gene for each gonadal tissue type separately. We used the normalized values on the same scale to test for sex differences at the adult stage.

Two additional datasets had stage-specific (i.e., tissues or cell types) expression level estimates in single cells for both somatic and germline cells from gonads for adult females (Zhang et al. 2018) and adult males (Guo et al. 2018). Pseudobulking the single cell expression data allowed us to test for the effect of adult male germline cell expression on the germline mutation rate without the noise introduced by somatic expression in the testis. Specifically, the adult human testis atlas (Guo et al. 2018) includes eight germline stages of spermatogenesis distinguished by cell marker expression (original stages from (Guo et al. 2018) indicated with numbers): spermatogonial stem cells (stage 1), differentiating spermatogonia (2), early primary spermatocytes (3), late primary spermatocytes (4), round spermatids (5), elongated spermatids (6), and finally, early (7) and later stages of sperm (8). The testis atlas also included five somatic testis tissues (i.e., Leydig cells, macrophages, Sertoli cells, endothelial cells, myoid cells). Using these data, we obtained a general measure of adult male germline transcription by pseudobulking the UMI counts from single-cell expression levels for the 8 stages of adult spermatogenesis (Guo et al. 2018) present within an individual donor for each of the 3 adult male donors. We normalized gene-specific UMI counts by dividing by the donor-specific total read count and by converting to CPM by multiplying by one million, incrementing by 1, and log transforming (base = 2). We averaged the donor-specific log_2_CPM values to obtain a single measure of expression per gene, our proxy for adult male germline expression level.

In the adult testis atlas, we could also consider the separate effects of each of the adult male somatic and germline stages on the total male mutation rate. To this end, we first pseudobulked the UMI counts per gene for each of the 3 donor x 13 stage combinations. To adjust for differences in cell counts across the 39 pseudobulked groups, we divided the pseudobulked counts by the total number of cells for each donor x stage group. We then adjusted for differences in sequencing depth by dividing the cell-averaged, gene-specific UMIs by the total number of UMIs present in the relevant donor, enabling us to obtain a measure of expression that could be compared among stages. The donor- and gene-specific values were converted to log_2_(CPM+1). Finally, we took the average of the expression values among the three donors to obtain one measure of expression per gene in each stage.

The female expression data (Zhang et al. 2018) were collected from five follicle stages, each consisting of an oocyte and the surrounding granulosa cells (with the temporal order of follicle maturation proceeding as primordial, primary, secondary, antral and preovulatory as identified in (Zhang et al. 2018)). Cell types were distinguished morphologically and through marker expression. Eight donors contributed unequally to the total sample of oocytes and granulosa cells (Zhang et al. 2018). To analyze these data, we first converted the expression units from FPKM to TPM, which provides relative transcript abundance and is more suitable for sample comparisons (Li et al. 2010; Pachter 2011; Wagner et al. 2012). TPM and CPM are both relative measures of expression that are appropriate to compositional data (Robinson et al. 2010; Friedman and Alm 2012; Fernandes et al. 2014; Love et al. 2014; Satija et al. 2015)). Because the primordial follicle stage can remain dormant for decades, the effects of expression in this cell type are important to consider separately. We therefore focused on two groups, one consisting of only primordial stage cells and one of all five stages together. To do so, we first calculated the average TPM expression for each cell type by donor combination. Then we took an average across donors for only primordial stage oocytes and a separate average across donors for all five oocyte stages together. We took the log (base=2) for the primordial-only and the all-stages datasets, resulting in one log_2_TPM value per gene for each group. The two measurements were highly correlated (ρ=0.91, P < 2.2x10^-16^).

### Additional covariates

We quantified the effect of expression levels on mutation rates by building a series of multiple regression models to estimate the effect of a predictor while holding all other predictors constant and controlling for confounding effects. All covariates were re-scaled (mean=0, standard deviation=1) in the models for interpretability. All regression models used individual genes as the data points.

To assess the effect of transcription on germline mutation rates, we used gene-specific expression data from one of the four datasets described above. Two datasets used the same measures of expression levels in the two sexes, enabling a joint comparison of the relationship in males versus females. The joint-sex models appropriately control for sex within their models; we also applied one FDR-correction on the p-values for the coefficients of all joint-sex models (Supplementary Tables 1, 2, and 7-11,).

We used the two other expression datasets to examine the effect of germline expression versus gonadal somatic expression in each sex separately. The single-sex models are used to confirm and support the results found in the joint-sex models. We applied an FDR-correction on the coefficient p-values for all of the male-specific regression models (Table 1 and Supplementary Tables 3,6) and a separate FDR-correction for the female-specific regression models (Table 2 and Supplementary Tables 4,5,13,14).

**Table 1.**
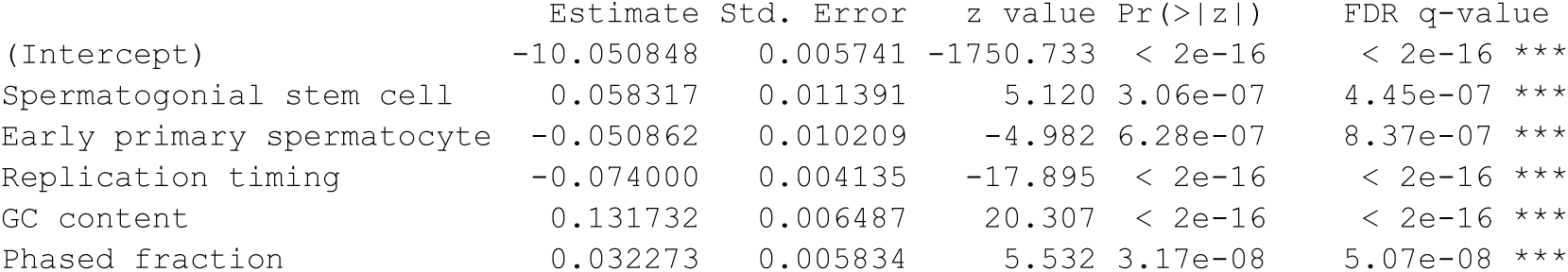
The effect of gene expression from adult testis cell types on male germline mutations using a Poisson multiple regression (for 15,220 genes). Non-significant cell stages were dropped. All predictors are re-scaled; a logged offset term is included to account for mutational opportunities per gene (i.e., gene length).

|  | Estimate | Std. Error | z value | Pr(>\|z\|) | FDR q-value |
| --- | --- | --- | --- | --- | --- |
| (Intercept) | -10.050848 | 0.005741 | -1750.733 | < 2e-16 | < 2e-16 *** |
| Spermatogonial stem cell | 0.058317 | 0.011391 | 5.120 | 3.06e-07 | 4.45e-07 *** |
| Early primary spermatocyte | -0.050862 | 0.010209 | -4.982 | 6.28e-07 | 8.37e-07 *** |
| Replication timing | -0.074000 | 0.004135 | -17.895 | < 2e-16 | < 2e-16 *** |
| GC content | 0.131732 | 0.006487 | 20.307 | < 2e-16 | < 2e-16 *** |
| Phased fraction | 0.032273 | 0.005834 | 5.532 | 3.17e-08 | 5.07e-08 *** |

**Table 2.** The effect of averaged oocyte expression (across five stages of folliculogenesis in adult females) on female germline mutations using a negative binomial regression model (for 15,200 genes). All predictors are re-scaled; a logged offset term is included to account for mutational opportunities per gene (i.e., gene length).

|  | Estimate | Std. Error | z value | Pr(>\|z\|) | FDR q-value |
| --- | --- | --- | --- | --- | --- |
| (Intercept) | -11.44065 | 0.01334 | -857.495 | < 2e-16 | < 2e-16 *** |
| Adult oocyte expression | 0.08352 | 0.01543 | 5.411 | 6.26e-08 | 8.57e-08 *** |
| Replication timing | -0.12161 | 0.01131 | -10.755 | < 2e-16 | < 2e-16 *** |
| GC content | 0.17268 | 0.01529 | 11.295 | < 2e-16 | < 2e-16 *** |
| Phased fraction | 0.04161 | 0.01387 | 3.001 | 0.00269 | 0.00318 ** |

Regression models of the effect of expression levels on the mutation rate included three covariates: GC content, replication timing, and the fraction of mutations that were phased to male or female genomes. GC content was included because it is known to affect the mutation rate, both directly (Michaelson et al. 2012; Kiktev et al. 2018), and indirectly, via the sequencing depth (Risso et al. 2011; Vallejos et al. 2017). Within each gene, we calculated GC percent for only the regions passing the 10X filter. Replication timing is also known to affect the mutation rate (Stamatoyannopoulos et al. 2009; Koren et al. 2012; Liu and Zhang 2020). We averaged the replication timing values of 300 iPSC lines (Ding et al. 2021) at the provided genomic locations (zero-scaled, i.e., with negative values denoting regions that replicate later than average and positive values earlier than average). We then averaged the positions that fell along the transcribed regions passing the 10X filter to obtain one replication timing value per gene. Of the 15,609 protein-coding genes present across all four expression data sets, 15,220 had replication timing information available.

When using read-backed phased mutations, we included the fraction of phased mutations as a covariate. A complication of using read-backed phasing is that we expect more mutations to be phased in regions with greater nucleotide diversity. Since genes with higher levels of transcription tend to be under stronger purifying selection (Zhang and Yang 2015), and thus have less diversity, the rate of phasing may itself depend on expression. Accordingly, we observed that the fraction of phased mutations (maternal and paternal) declines with increasing male germline expression across deciles. This negative correlation is potentially problematic because the sex-specific mutation rate may appear to decline with expression simply because of its correlation with phasing rate. To take this confounding effect into account, we included the fraction of mutations that were phased per decile as a covariate when regressing mutation rates on expression levels. However, the phased fraction is imprecisely estimated per gene. We therefore assigned each gene a phased fraction value by using the value from the expression decile to which it belonged. We calculated this phased fraction by summing the number of all male and female phased mutations within a given expression decile and dividing by the total number of mutations in that decile. Phased fraction was not included as a covariate for models using only mutations that were phased by third generation transmission, since virtually all mutations are phased by this approach, or for models using only sperm and seminiferous tubule mutations, which are of male origin.

Since CpG transitions occur at a greater rate when the C is methylated, we included methylation level as a covariate in the regression models when CpG>TpG mutation rate was the response. We used methylation data from Gene Expression Omnibus for ovaries (GSM1010980) and testis (GSE127321, GSE127281) (ENCODE Project Consortium 2012). Average methylation values for each gene were calculated by averaging the methylation values for all CpGs in the 10X filtered regions. This calculation was performed in the ovary and testis datasets separately. The regressions excluded genes without CpG dinucleotides or genes without any methylation data. To avoid biasing our estimates, genes with zero CpG>TpG mutations were retained so long as CpG methylation data were available.

### Modeling approach

We implemented a Poisson GLM multiple regression model (R command glm(family=poisson(link=log)) with the number of mutations per gene as the response variable, with the number of surveyed base pairs as an offset term to account for gene length (i.e., denominator). If the data suggested mild overdispersion (dispersion parameter between 1.1 and 1.4) with respect to a Poisson model, we applied the next most simple quasipoisson model (Crawley 2013). In the case of more severe overdispersion (>1.4), we opted to use a negative binomial model (Crawley 2013). In general, we detected low or acceptable levels of collinearity across the covariates in the regression models (VIF < 5 or GVIF < 3) (Fox and Monette 1992). For greater levels of collinearity, we checked whether GLM coefficients were stable by comparison to LASSO and ridge regression coefficients. In particular, for the male stage-specific regression model, which included expression from multiple somatic and germline cell stages, we conducted a backwards elimination that dropped non-significant stages to only a pared down set with a significant effect on the mutation rate. To assess the stability of the remaining coefficients, we assessed whether LASSO and ridge regressions yielded similar values.

### Male-mutation bias

To test whether the ratio of the male-to-female mutation rates (α) differs significantly between any two quantiles of expression level, we bootstrap resampled the male and female mutation rates of each gene according to the sex-specific quantile 10,000 times. For each of the 16 male by female quantile combinations (Figure 3), we then calculated the ratio of the male-to-female mutation rates, yielding a distribution of 10,000 α ratios per cell. Next, we computed the difference of the ratios for the smallest and highest valued cells (e.g., male quantile 1 and female quantile 3 versus male quantile 3 and female quantile 1 from Figure 3). We examined whether the 95% confidence interval for the bootstrapped difference excluded zero for these two cells of interest.

### Asymmetry measure

By choosing the frame of reference (e.g., mutation type or strand), it is possible to separate changes of paired bases and tally the changes on each strand individually. For a given mutation type, T-asymmetry is measured by the difference in the number of base changes on the transcribed versus non-transcribed strands. For example, if the transcribed strand is 5’AAATT in both parents and 5’AAGTC in the offspring, one third of the A bases on the transcribed strand were changed to G; one half of the A bases were changed to G on the non-transcribed strand. Picking a reference base (e.g., A instead of T) to count on each strand is arbitrary. We analyzed strand asymmetry in seven basic mutation types: A>G, A>C, A>T, C>A, C>G, nonCpG C>T, and CpC>TpG.

We tested for differences in T-asymmetry through a proportions test (chi-square test) by counting the total number of DNMs of a given type on the non-transcribed strand (e.g., A>G) versus the total number of mutations of that same type on the transcribed strand. To account for mutational opportunities, we counted the total number of relevant nucleotides on the non-transcribed strand (i.e., A) versus the total number of the same nucleotide on the transcribed strand. We estimated the degree of strand asymmetry in the same set of genes for which we have data in males and females for all four expression datasets and the replication timing dataset. We plotted the log_2_ (non-transcribed strand mutation rate / transcribed strand mutation rate) and used the delta method to calculate 95% confidence intervals on the T-asymmetry estimates to evaluate significance.

We compared the extent of asymmetry between the sexes by performing a two-tailed, Z-test on the difference of the log_2_ (non-transcribed / transcribed strand) for a given type of mutation; P-values for tests of sex differences were FDR-adjusted.

## Results

### Joint models for the effect of expression levels

To analyze sex differences in the effect of transcription on mutagenesis in the germline, we used fetal single-cell germline expression from the Reproductive Cell Atlas (Garcia-Alonso et al. 2022) and adult bulk testis and ovary tissues from GTEx (GTEx Consortium 2013) (see Figure 1). In two separate models, we regressed the rate of DNMs on the fetal or adult gonadal expression levels for the male and female germlines jointly. Both models included sex as a predictor and the covariates of GC content, replication timing, and phased fraction. We considered 15,220 protein-coding genes and included only genes for which we had expression level estimates in both sexes and replication timing values (see *Materials and Methods*). We used 47,787 paternal and 12,712 maternal germline DNMs identified from pedigree sequencing, which were phased using other variants in reads or third generation transmission (Goldmann et al. 2016; An et al. 2018; Halldorsson et al. 2019). The mutation rate was calculated as the number of DNMs in a transcribed region, adjusted by the length of the region, after removing sites with potentially low sequencing depth using the 10X filter requirement. If we instead applied no filter on coverage or a more stringent 20X filter on both mutations and transcribed regions, we obtained the same qualitative results.

Using the Reproductive Cell Atlas dataset, we found that males and females differ significantly for the effect of fetal germline expression on the mutation rate (Figure 2 and Supplementary Table 1). Expression levels have no discernible effect on the mutation rate in males (P=0.50; Supplementary Table 1), but a positive effect in females (P=1.7x10^-8^); the female slope differs significantly from the male slope (P = 4.02x10^-6^; Supplementary Table 1; Figure 2). Similarly, using the GTEx data set, we found that bulk adult testis tissue expression levels have no detectable effect on the male germline mutation rate (P=0.09; Supplementary Table 2), but that bulk adult ovary tissue expression levels have a positive effect on the female germline mutation rate (P = 3.7x10^-9^); the slopes again differ significantly between the sexes (P = 0.001; Supplementary Table 2). Finally, including sex as a predictor explains significantly more variation in the final model than a model without it for both the fetal (F-test: P < 2.2x10^−16^) and adult gonadal expression (F-test: P < 2.2x10^-16^). Both fetal germline expression and adult gonadal expression suggest that there is no discernible effect of expression levels on mutation rates in males but a positive effect in females; in other words, the relationship between expression level and mutation rate depends on sex. As expected, GC content and phased fraction are positively associated with the mutation rate, and replication timing is negatively associated with mutations in both of the joint-sex regression models (Supplementary Tables 1 and 2).

**Figure 2.**
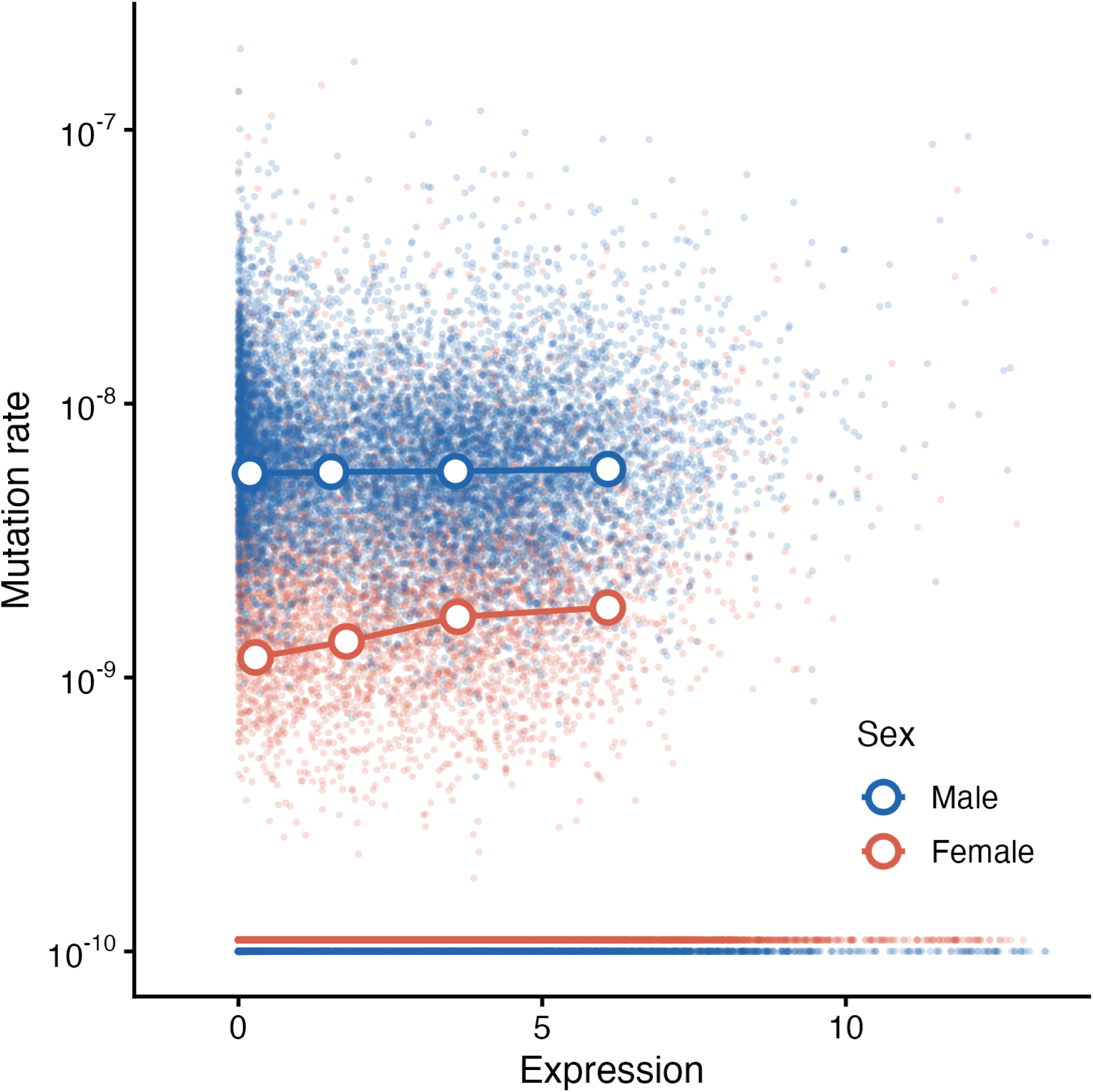
Scatterplot of sex-specific mutation rates as a function of pseudobulked fetal expression levels. Mutation rates per genome (averaged across 7,596 trios) are plotted for each of 15,220 genes (filled-in points) on a log_10_ scale, and expression is in log_2_CPM. Genes with zero mutations are shown along the bottom of the plot, at an arbitrary value of 10^-10^. Circles plot the averaged model-predicted mutation rates based on Supplementary Table 1 for expression level quantiles (determined separately in males and females); these corrected mutation rates were adjusted for the covariates of GC content, replication timing, phased fraction and interactions by subtracting their mean contributions.

**Figure 3.**
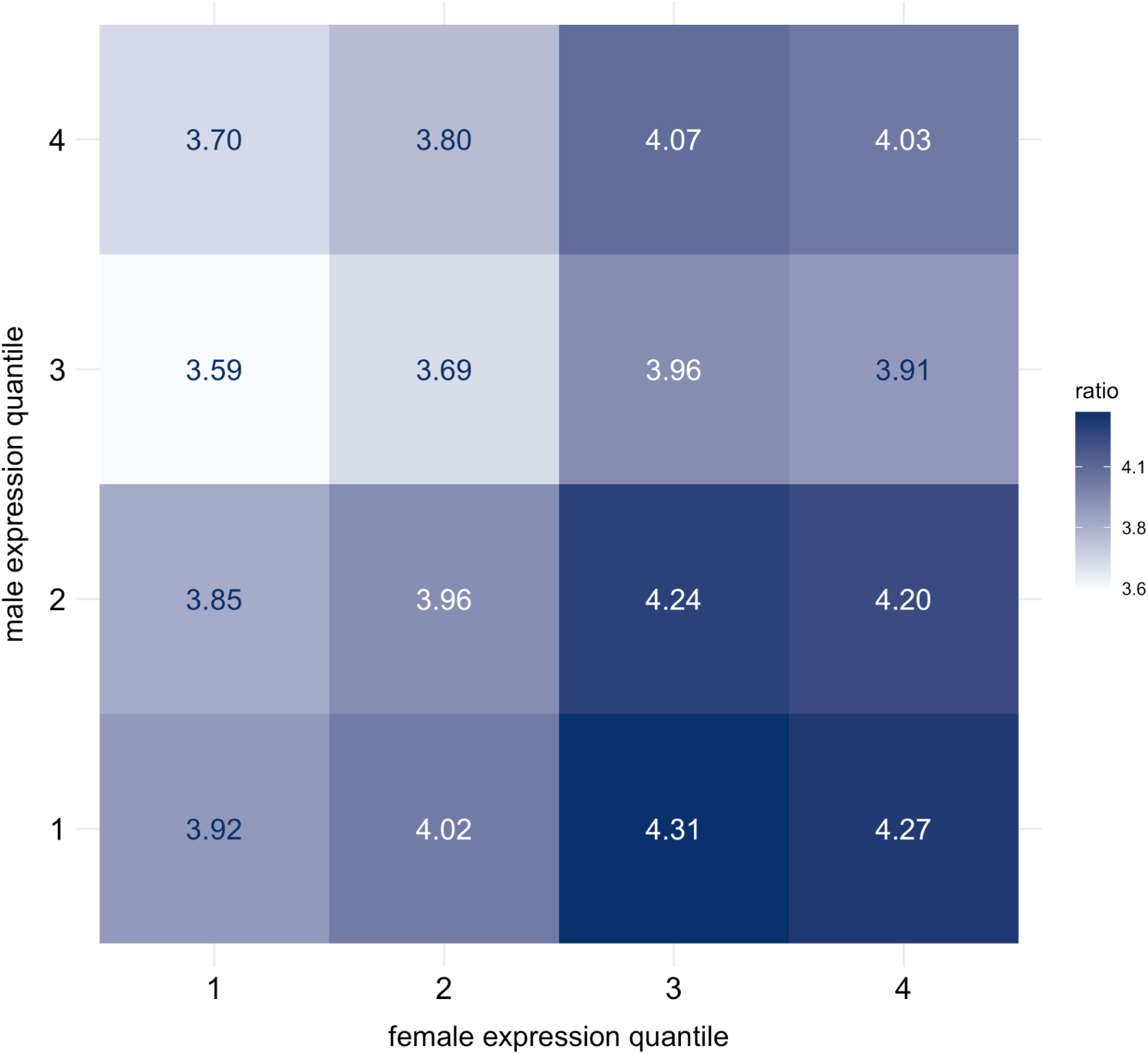
The ratio α of paternal to maternal mutations for different expression levels in males and females. In each sex, genes were assigned to four quantiles using expression levels from the pseudobulked fetal germline expression data. The absolute difference between the largest ratio (male quantile 1 and female quantile 3, α=4.31) and smallest ratio (male quantile 3 and female quantile 1, α=3.59) equals 0.72; the bootstrapped 95% confidence interval of this difference excluded zero [0.1082, 1.3071] (see Materials and Methods).

The effects of covariates are stable across datasets. In the two joint-sex models, the coefficients for GC content (β ≈ 0.15, SE = 0.007) and replication timing (β ≈ -0.1, SE = 0.004) are similar, suggesting that their effects are robust and not sensitive to the expression data used. The coefficients for male phased fraction are comparable (β ≈ 0.3) but statistically significant for only the model using the GTEx data. The coefficients for female phased fraction are significant and consistent in both data sets: the 95% confidence interval is [0.14, 0.26] in the fetal expression data (Supplementary Table 1) versus [0.17, 0.26] in the GTEx data (Supplementary Table 2). The estimate of the female-specific expression effect was somewhat larger in the fetal germline expression data (β = 0.19 and 95% CI = 0.13-0.26; Supplementary Table 1) than in the adult gonadal expression data (β = 0.11 and 95% CI = [0.07, 0.14] from Supplementary Table 2), but the 95% confidence intervals overlapped. The coefficient of scaled female expression on the female germline mutation rate is similar in magnitude to the scaled coefficients for GC content and replication timing, suggesting that female expression is comparable in influence to well-established mutational covariates.

### Individual sex models for the effect of expression levels

The Reproductive Cell Atlas and GTEx datasets enable a direct comparison between males and females, but both have limitations. For one, fetal expression levels may not be pertinent to germline mutations that accrue during adulthood. Moreover, somatic cells included in the GTEx gonadal data contribute noise that may mask effects of germline transcription. For these reasons, we chose to use two additional datasets from adult germline cells to confirm relationships of expression levels and mutation rates inferred in the joint-sex regression models. Because these additional datasets came from separate studies (Guo et al. 2018; Zhang et al. 2018) and employed different measurement scales and techniques, we could not directly compare the sexes and instead constructed separate sex-specific regression models.

For males, we obtained a proxy for overall adult male germline expression by pseudobulking single-cell RNAseq expression from 8 cell stages of spermatogenesis (see *Materials and Methods*). This pseudobulked expression is akin to GTEx bulk testis tissue expression but does not include somatic cell transcription. Using this proxy for adult male germline expression, we confirmed the lack of an effect of expression levels on the male germline mutation rate after adjusting for the covariates of GC content, replication timing, and phased fraction (P=0.65; Supplementary Table 3). This confirmation is expected, given the high positive correlation between the adult and fetal expression values (ρ = 0.67; P < 2.2x10^-16^). Regardless, these analyses indicate that neither the fetal nor the adult germline expression levels are associated with the male germline mutation rate.

For females, we constructed a multiple regression model that used the average expression of the five oocyte stages as a proxy for adult female germline transcription (see *Materials and Methods*). By this approach, there is a significantly positive effect of oocyte expression levels (P = 6.3x10^-8^; Table 2) on the female germline mutation rate after adjusting for the covariates of GC content, replication timing, and phased fraction. The averaged adult oocyte expression levels are highly correlated with female fetal expression values (ρ = 0.74, P < 2.2x10^-16^). Furthermore, using only averaged expression levels from primordial oocytes, the longest lasting stage of folliculogenesis, also shows a positive effect on the female germline mutation rate using a negative binomial model (P < 5.7x10^-9^; Supplementary Table 4). We note that the parental ages of the female (mean = 28y) and male (mean = 21y) donors for the adult germline expression data were not different (t-test, P=0.1) and thus not likely to explain the sex differences in the associations with adult germline expression.

### Consistency checks that do not rely on read-based phasing

To verify the finding of a null association in males and a positive association in females, we conducted a number of checks. First, as an alternative to including the phasing rate as a covariate, we limited the analysis to the (much smaller) subset of mutations that were phased using transmission to a third generation (4,865 paternal and 1,136 maternal mutations) (Jónsson et al. 2017), an approach that does not rely on the presence of nearby SNPs. We again found that male germline expression levels have no effect on male mutation rates, whether using adult gonadal (P=0.11) or fetal germline expression (P=0.58). In females, expression levels have a positive effect on such mutations whether using adult germline (P=0.04) or fetal germline expression (P=0.03).

As a second approach, we considered the total number of mutations available in our filtered regions as our response variable, whether or not they were phased and regardless of parental origin. We note that ∼80% of these 194,356 mutations are expected to have occurred in the male germline (Jónsson et al. 2017). Owing to this predominance, if there is truly no effect of male germline expression levels, we expect no association with male expression when considering all mutations (phased or not). In turn, if only female expression levels have an effect on female germline mutations, the male mutations should contribute noise but not distort an underlying positive relationship. After performing this analysis, we recovered a positive association between the total mutation rate and female adult germline expression levels, as measured by the average of all oocyte types (P < 2x10^-16^; Supplementary Table 5), or by only primordial oocytes (P = 2x10^-16^). We observed no significant relationship with male adult germline expression levels as measured by the pseudobulked single-cell expression values (P=0.42; Supplementary Table 6). Similarly, using all available mutations, we constructed a model that included GC content, replication timing, and both male and female fetal germline expression data as covariates, as well as male and female phasing rates. This model indicates an effect of female, but not male, expression (Supplementary Table 7).

As a final test, we considered only mutations with known male origin, namely those identified in sperm (Abascal et al. 2021; Neville et al. 2025; Shoag et al. 2025) and seminiferous tubules (Moore et al. 2021). Using these 14,482 mutations, we find no effect of the male germline expression from pseudobulk-derived adult (P=0.998) or fetal germline expression values (P=0.22) on the male germline mutation rate, after controlling for GC content and replication timing. These additional analyses therefore confirm that there is no or only a very weak association of male expression levels and paternal germline mutation rates.

### Paternal mutational bias in protein-coding regions

To date, estimates of paternal bias, whether from DNM calls in pedigrees or comparisons of X chromosome and autosomal divergence rate have mainly been genome-wide (e.g., (Wilson Sayres and Makova 2011; Jónsson et al. 2017; de Manuel et al. 2022)). However, the finding of a positive association between mutation and expression in females and a null association in males suggests that at least in protein-coding genes, the paternal bias depends upon expression levels (as well as associated factors such as GC content and replication timing). Indeed, the observed data (not adjusted for covariates) indicate that the male mutation bias can differ by as much as 20% when partitioning genes by four expression level quantiles across both males and females (Figure 3). Therefore, the extent of paternal bias in a species is not a fixed quantity, but can vary substantially by genome context.

### Effects for CpG sites versus other mutation types

Next, we sought to understand whether the sex difference only pertains to a subset of mutations. Previous analyses have analyzed germline mutations in terms of COSMIC single base substitution (SBS) mutational signatures identified in tumors (Rahbari et al. 2016, Moore et al. 2021; Spisak et al. 2024), with each signature defined as a distribution of mutation counts over 96 SBS types (Alexandrov et al. 2013; Alexandrov et al. 2020); (Koh et al. 2021). In both males and females, most germline mutations are assigned to SBS5 and to a lesser extent SBS1 (Rahbari et al. 2016; Moore et al. 2021). Stratified by expression levels, however, there are too few DNMs for us to analyze and thus a concern that the signature assignment would be unreliable (Serrano Colome et al. 2023). Instead, we divided germline mutations into two sets: transitions at CpG sites, which constitute the bulk of SBS1 mutations (Alexandrov et al. 2013) and are thought to arise from deamination of methylated CpGs (Guo et al. 2022) or errors in replication (Seplyarskiy et al. 2021; Tomkova et al. 2024), and all other mutation types, most of which likely contribute to the relatively flat SBS5 signature, whose etiology is unknown but may reflect the error profile of translesion synthesis and repair (Spisak et al. 2025).

We first regressed all mutation types except transitions at CpGs on the expression levels for the male and female germlines jointly. The multiple regression model included the covariates of sex, phasing rate, GC content, and replication timing. Using GTEx data, the two sexes differ significantly in the relationship of transcription levels and sex-specific mutation rates (P = 0.0002; Supplementary Table 8), with a positive slope for female mutations (P=7.6x10^-9^) and no discernible relationship for male mutations (P=0.06). The same patterns were found using fetal germline expression values from the Reproductive Cell Atlas (Supplementary Table 9); the sexes differ significantly (P= 6.5x10^-8^), and the effect of expression levels is positive in females (P=6.8x10^-12^) and null in males (P=0.42). Thus, when we consider all mutations other than CpG transitions, they recapitulate what is observed for all mutations, as expected.

Next, we focused on the subset of mutations involving transitions at CpG dinucleotides and regressed their mutation rate on the GTEx expression levels for the male and female germlines jointly. The multiple regression model included the covariates of sex, phasing rate, GC content, and replication timing. Because mutation rates are substantially higher for the subset of CpG dinucleotides that are methylated, we included methylation levels derived from adult ovaries and testes as a covariate; these methylation data are from the same tissue type as was used for the GTEx data. Surprisingly, the mutation rate of CpG transitions is negatively correlated with expression levels (P=0.01; Supplementary Table 10), similarly in both sexes (testing for a difference, P=0.78). Both the overall negative effect of expression levels (P=0.02) and the lack of sex difference (P=0.39) are also seen when using fetal germline data from the Reproductive Cell Atlas (Supplementary Table 11).

### Stage-specific gene expression levels

If transcription contributes to mutagenesis, it may do so differentially across cell stages of gametogenesis. We explored this possibility in males and females separately using expression levels from different germline and somatic cell types found in adult gonads. First, in males, we constructed a stage-specific regression model that included the comparable expression values for the eight germline and five somatic cell types from adult testes (Guo et al. 2018), and the three covariates of GC content, replication timing, and phased fraction, as well as a logged offset term to account for mutational opportunities. We note that while, in principle, it might be preferable to weight the contribution of individual stages by their duration (or according to the number of cell divisions within each stage), any such weighting would be highly imprecise, given the limited developmental timing information on spermatogenesis in humans and the variation in parental ages.

We performed a backwards elimination on the full stage-specific model, progressively dropping cell types that had no significant effect on the male mutation rate (i.e., P > 0.05). This stepwise procedure removed all five somatic stages, as expected. The resulting model retained two spermatogenic cell stages (numbers indicate position in the temporal order): spermatogonial stem cells (stage 1) and early primary spermatocyte (3). Male mutation rates are positively associated with gene expression levels in the spermatogonial stem cell stage after controlling for expression level from the primary spermatocyte stage. Conversely, male mutation rates are negatively associated with expression levels from the primary spermatocyte stage after controlling for expression levels from the stem cell stage (Table 1). As expected, GC content and phased fraction were positively associated with the mutation rate, and replication timing was negatively associated (Table 1).

Changes in the sign of the slopes across temporally ordered cell stages could indicate that the coefficient estimation was unstable due to multicollinearity among the cell stages. To explore this possibility, we also performed LASSO and ridge regressions, two regularization approaches that reduce overfitting and the effect of predictor correlations. We obtained coefficients with similar values as in the Poisson regression (Supplementary Table 12), suggesting that the estimated coefficients are robust. The retention of these two stages, as well as the opposite signs of their coefficients, suggests that a measure of expression that averages over several germline stages may mask the nuanced effects of transcription on mutations.

Second, in females, we constructed a stage-specific regression model that included the averaged oocyte and averaged granulosa cell expression levels from adult females (Zhang et al. 2018), the three covariates of GC content, replication timing, and phased fraction, and a logged offset term for the mutational opportunity. The mutation rate increases significantly with the averaged oocyte expression levels (Table 2 and Supplementary Table 13). As expected, GC content and phased fraction are positively associated with the mutation rate and replication timing is negatively associated (Table 2 and Supplementary Table 13).

Curiously, given that somatic cells should not influence germline mutation rates, the average expression across five adult granulosa cell stages is negatively associated with the female germline mutation rate in the full stage-specific model (Supplementary Table 13). When only one of the two cell types is included in a model with the other covariates, oocyte expression alone is a significant positive predictor (Table 2) whereas granulosa cell expression alone is not (Supplementary Table 14; P = 0.26). Moreover, the effect of oocyte expression levels becomes larger when granulosa cell expression levels are included in the model (β = 0.08 alone vs. β = 0.14 in the full model). In this regard, we note that oocyte and granulosa expression are highly correlated (r = 0.69, P < 2.2×10⁻¹⁶). Together these patterns suggest that oocyte expression levels lie on the causal path to female germline mutation rates, whereas the effect of granulosa cell expression levels reflects their role as a suppressor variable (Friedman and Wall 2005).

### Sex-specific T-asymmetry

T-asymmetry arises because the balance of repair (TCR) and damage (TCD) differs between the transcribed and non-transcribed strands. A negative association between expression levels and mutation rate, together with appreciable T-asymmetry, indicates that TCR has a larger net effect than TCD. By contrast, a positive association combined with strong T-asymmetry may indicate that the effects of TCD dominate. Finally, the lack of an association between expression levels and mutation rate, coupled with the absence of T-asymmetry, suggests the lack of both TCR and TCD, or that the two effects cancel out.

Several mutation types demonstrate significant T-asymmetry in one or both sexes (Figure 4). Consistent with previous reports (Seplyarskiy and Sunyaev 2021) using a subset (Jónsson et al. 2017) of the available germline data, A>G and C>G asymmetry is stronger in females than in males. We also found stronger T-asymmetry for A>C, nonCpG C>T, and CpG > TpG in females.

**Figure 4.**
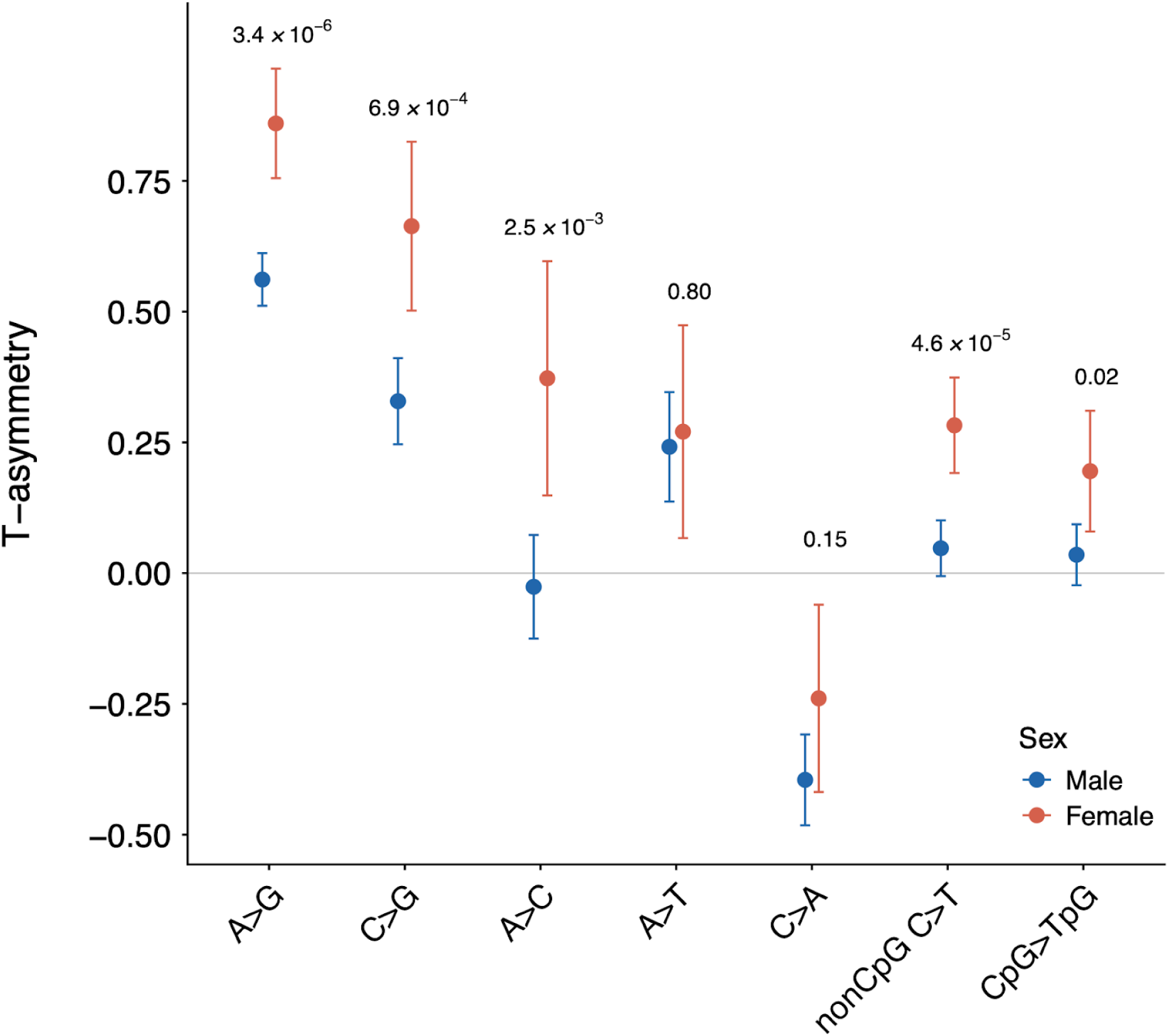
T-asymmetry across mutation types, calculated as log_2_ of the ratio of the mutation rates on the non-transcribed to transcribed strands. Whiskers denote the 95% confidence intervals. FDR adjusted P-values are provided above a pair of bars for a two-tailed test of a sex difference in the T-asymmetry values.

## Discussion

In males, we found no effect of transcription levels on germline mutations when all mutation types (CpG transitions or not) are combined: there is no detectable effect when using all paternally phased mutations, the subset of paternal mutations identified through third generation transmission, or only mutations in seminiferous tubules and sperm. We also detected no association whether relying on adult pseudobulked single-cell, fetal pseudobulked single-cell, or adult bulk testes tissue expression data. The lack of an effect of transcription on mutation rates in males suggests either that transcription-related damage and repair have no or very weak effects throughout male gametogenesis or that, summed over all stages of spermatogenesis, the effects of repair approximately cancel out that of transcription-related damage.

In support of the latter possibility, expression levels in a subset of spermatogenic stages are associated with the male mutation rate (Table 1): positively in spermatogonial stem cells and negatively associated in primary spermatocytes. Consistent with significant associations across individual stages, there is significant T-asymmetry for A>G, A>T, C>A and C>G in the male germline (Figure 4). A caveat is that T-asymmetry and replication associated asymmetry (R-asymmetry) are correlated in humans (Seplyarskiy et al. 2019; Koyanagi et al. 2022), so that the T-asymmetry could, at least in part, actually be indicative of R-asymmetry. That said, recent empirical observations in a murine model indicate that TCR is a discernible, and much stronger, source of strand bias compared to replication (Anderson et al. 2024). Therefore, the male T-asymmetry observed here could plausibly have emerged from individual germline stages experiencing net mutagenic or net reparative transcription, despite an overall null association with pseudobulked fetal germline, pseudobulked adult germline and bulk testis tissue expression. Alternatively, while this study found no effect of total steady-state RNA, it may be that nascent transcript levels are better for measuring the effect of expression on mutations in males and for potentially explaining the presence of T-asymmetry (Anderson et al. 2024).

In females, by contrast, mutation rates increase with expression levels, whether considering all maternally phased mutations or only the subset of maternal mutations identified from third generation transmission. Primordial oocytes have relatively less transcription (Zhang et al. 2018), suggesting that DNA repair may also be less active and perhaps less able to redress damage arising from the limited transcription present in the fetal germline and in adult primordial oocytes. Interestingly, in the male germline, the transcriptionally quiescent spermatogonial stem cell stage (Guo et al. 2018) also demonstrated a positive association with mutation. Downregulation of repair pathways due to decreased global expression may partially explain these positive associations in males and females, but further human-specific empirical data is required to support this speculation.

Regardless of the precise mechanism, it appears that germline transcription in females inflicts more damage than is efficiently and accurately repaired. The positive association in females is present in the pseudobulked fetal germline, the average expression of all adult oocyte stages and in bulk ovary tissue expression. The consistent pattern over development may explain why T-asymmetry is more pronounced in females than males (Figure 4).

While our findings point to a difference between the sexes in the impact of overall transcription levels on mutagenesis, the difference may also be attributable to sex-specific covariates that we do not fully capture. For instance, we use replication timing data from iPSC lines rather than from sex-specific germline cells, data which, to our knowledge, do not exist for the human germline. Thus, the causal path that gives rise to the sex difference revealed by our analysis remains to be further elucidated. Moreover, we examined effects at the level of the entire gene, when there may be more local effects of expression on mutation rates, e.g., around transcription start sites (Cortés Guzmán et al. 2025), which remain to be explored using a denser set of DNMs.

Nevertheless, the general increase in mutations with expression in females but not males contrasts with what one might expect from the overall paternal bias in germline mutations (Jónsson et al. 2017).

Our results further indicate that the male-mutation bias in protein-coding regions should not be viewed as a fixed species-specific parameter. Rather, the male-mutation bias, which differed up to ∼20% in our study, depends on sex-specific expression levels and associated covariates. Hence, the variation in the extent of male-mutation bias in the protein-coding regions of other species will depend on the exact sex-specific correlations between expression levels and mutation rates and the associated genomic variables. One implication is that slight differences in the sex bias of different mammals could reflect changes in transcription levels (or covariates) during gametogenesis.

Curiously, we observed a negative association between the rate of CpG transitions and expression levels that was not sex-specific and that was supported by the presence of T-asymmetry (Figure 4). A similar negative association has recently been reported for somatic SBS1 mutations, albeit without covariate adjustment (Silveira et al. 2024), and with no T-asymmetry (Alexandrov et al. 2020; Silveira et al. 2024). A negative association between expression and CpG>TpG mutations suggests the possibility that transcription can trigger the repair of deaminated methylated cytosines, a BER-specific lesion, through TCR itself or by TCR-mediated BER recruitment (Svejstrup 2002; Kim and Jinks-Robertson 2010; Chakraborty et al. 2021). Alternatively, the association between expression levels and CpG>TpG mutation rates may not be causal. In other words, the models may be lacking an important confounder for this mutation type in both the adult male gonadal and fetal male germline expression analyses. If so, the low levels of T-asymmetry at CpG>TpG mutations in males may reflect the small subset of mCpGs experiencing transitions due to mutational mechanisms that do not come from spontaneous deamination (e.g., as part of signature SBS5).

Nevertheless, the finding that the overall relationship of expression levels and mutation rates differs between the sexes, as well as among the individual stages within males, highlights that the germline mutation rate results from effects within multiple distinct cell types. Further insight will be gained by studying effects of damage and repair for each cell type separately, rather than by making inferences from their cumulative effects.

## Data availability

This paper relied on publicly available data from previous publications (see Materials and Methods for references). Data tables and accompanying analyses are available through https://github.com/minyoungWyman/germExpressMuts.

## Acknowledgments

We thank S. Sanders for sharing unpublished phased mutational data, as well as Z. Yan and J. Guo for sharing unpublished data. We are grateful to the members of the Przeworski, Sella, and Andolfatto labs, especially F. Wu, V. Getseva, A. Stolyarova for comments, and the High Performance Cluster Computing at Columbia University for computational support. We thank M.C. Wyman.

