## Supplementary materials for "Sex differences in transcription-associated mutagenesis in the human germline"

Minyoung J. Wyman[1,*]

Ipsita Agarwal[1,2]

Marc de Manuel[1,3]

Natanael Spisak[1, 4]

Molly Przeworski[1,5]

[1] Department of Biological Sciences, Columbia University, New York, NY 10027, USA

[2] current address: Department of Statistics, University of Oxford; Oxford OX1 3LB, United Kingdom

[3] current address: Institute of Evolutionary Biology (CSIC-UPF). Passeig Marítim de la Barceloneta 37-49; 08003 Barcelona, Spain

[4] current address: Institut Imagine, INSERM, Paris, France

[5] Department of Systems Biology, Columbia University, New York, NY 10027, USA

Supplementary Table 1. The effect of fetal germline expression levels from the Reproductive Cell Atlas on phased mutations. See Materials and Methods for the details of pseudobulked fetal expression values used in the quasipoisson regression model (dispersion parameter =1.3; n=15,220 genes). For the intercept and all sex-specific coefficients, the male estimate is shown first (baseline) and then the difference between the male and female coefficients ("female diff."). A model comparison supported retaining the two-way interaction between phased fraction and sex (F-test: P=0.001). The final model explains substantially more variance than a model without sex (F-test: P < 2.2x$10^{-16}$).

| | Estimate | Std. Error | t value | Pr(>\|t\|) | FDR q-value | |
|---|---|---|---|---|---|---|
| (Male Intercept) | -10.056780 | 0.006414 | -1567.917 | < 2e-16 | < 2e-16 | *** |
| Fetal male germ. express. | 0.012872 | 0.018965 | 0.679 | 0.4973 | 0.4973 | |
| Intercept (female diff.) | -1.322150 | 0.011749 | -112.531 | < 2e-16 | < 2e-16 | *** |
| Male phased fraction | 0.033503 | 0.018305 | 1.830 | 0.0672 | 0.0798 | |
| Replication timing | -0.099071 | 0.004368 | -22.680 | < 2e-16 | < 2e-16 | *** |
| GC content | 0.149886 | 0.006576 | 22.793 | < 2e-16 | < 2e-16 | *** |
| Fetal germ. express. (female diff.) | 0.180933 | 0.039239 | 4.611 | 4.02e-06 | 6.74e-06 | *** |
| Phased fraction (female diff.) | 0.166290 | 0.033887 | 4.907 | 9.29e-07 | 1.61e-06 | *** |

Supplementary Table 2. The effect of GTEx gonadal expression levels on phased mutations. See Materials and Methods for details of the testis and ovary expression data used for the quasipoisson regression model (dispersion parameter =1.3; n=15,220 genes). For the intercept and all sex-specific coefficients, the male estimate is shown first (baseline) and then the difference between the male and female coefficients ("female diff."). A model comparison supported retaining the two-way interactions between phased fraction and sex (F-test: P=2.9x$10^{-13}$). The final model explains substantially more variance than a model without sex (F=test: P < 2.2x$10^{-16}$) (Male GTEx = testis bulk tissue expression; female GTEx = ovary bulk tissue expression).

| | Estimate | Std. Error | t value | Pr(>\|t\|) | FDR q-value | |
|---|---|---|---|---|---|---|
| (Male Intercept) | -10.055122 | 0.006569 | -1530.647 | < 2e-16 | < 2e-16 | *** |
| Male GTEx express. | 0.027240 | 0.015500 | 1.757 | 0.078863 | 0.09176 | |
| Intercept (female diff.) | -1.317691 | 0.011609 | -113.506 | < 2e-16 | < 2e-16 | *** |
| Male phased fraction | 0.036611 | 0.010665 | 3.433 | 0.000598 | 0.000874 | *** |
| Replication timing | -0.098866 | 0.004297 | -23.010 | < 2e-16 | < 2e-16 | *** |
| GC content | 0.154194 | 0.006499 | 23.725 | < 2e-16 | < 2e-16 | *** |
| GTEx express. (female diff.) | 0.086569 | 0.024662 | 3.510 | 0.000448 | 0.000672 | *** |
| Phased fraction (female diff.) | 0.186781 | 0.025317 | 7.378 | 1.65e-13 | 3.36e-13 | *** |

Supplementary Table 3. The effect of pseudobulked adult male germline expression on paternally phased germline mutations using a Poisson regression model (n=15,220 genes).

| | Estimate | Std. Error | z value | Pr(>\|z\|) | FDR q-value | |
|---|---|---|---|---|---|---|
| (Intercept) | -10.048374 | 0.005670 | -1772.260 | < 2e-16 | < 2e-16 | *** |
| Adult male germ. express. | 0.005195 | 0.011446 | 0.454 | 0.649905 | 0.6499 | |
| Replication timing | -0.076891 | 0.004075 | -18.869 | < 2e-16 | < 2e-16 | *** |
| GC content | 0.135183 | 0.006463 | 20.915 | < 2e-16 | < 2e-16 | *** |
| Phased fraction | 0.038131 | 0.010629 | 3.588 | 0.000334 | 0.000411 | *** |

Supplementary Table 4. The effect of averaged primordial oocyte expression (from adult females) on maternally phased germline mutations using a negative binomial model (n=15,220 genes).

| | Estimate | Std. Error | z value | Pr(>\|z\|) | FDR q-value | |
|---|---|---|---|---|---|---|
| (Intercept) | -11.43893 | 0.01334 | -857.584 | < 2e-16 | < 2e-16 | *** |
| Adult primordial oocyte expression | 0.08688 | 0.01492 | 5.825 | 5.72e-09 | 8.26e-09 | *** |
| Replication timing | -0.11825 | 0.01122 | -10.537 | < 2e-16 | < 2e-16 | *** |
| GC content | 0.16706 | 0.01531 | 10.914 | < 2e-16 | < 2e-16 | *** |
| Phased fraction | 0.03521 | 0.01317 | 2.673 | 0.00751 | 0.00814 | ** |

Supplementary Table 5. The effect of averaged oocyte expression (across five stages of folliculogenesis in adult females) on all available germline mutations, whether or not phased, using a quasipoisson regression model (dispersion parameter=1.3; n=194,356 mutations and 15,220 genes).

| | Estimate | Std. Error | t value | Pr(>\|t\|) | FDR q-value | |
|---|---|---|---|---|---|---|
| (Intercept) | -8.635841 | 0.003179 | -2716.291 | < 2e-16 | < 2e-16 | *** |
| Adult oocyte expression | 0.030588 | 0.003538 | 8.646 | < 2e-16 | < 2e-16 | *** |
| Replication timing | -0.047185 | 0.002265 | -20.828 | < 2e-16 | < 2e-16 | *** |
| GC content | 0.111293 | 0.003588 | 31.014 | < 2e-16 | < 2e-16 | *** |
| Phased fraction | 0.008179 | 0.003020 | 2.708 | 0.00677 | 0.00765 | ** |

Supplementary Table 6. The effect of pseudobulked adult male germline expression on all available germline mutations, whether or not phased, using a quasipoisson regression model (dispersion parameter=1.3; n=194,356 mutations and 15,220 genes).

| | Estimate | Std. Error | t value | Pr(>\|t\|) | FDR q-value | |
|---|---|---|---|---|---|---|
| (Intercept) | -8.636115 | 0.003199 | -2699.620 | < 2e-16 | < 2e-16 | *** |
| Adult male germ. express. | 0.005186 | 0.006376 | 0.813 | 0.4160 | 0.4437 | |
| Replication timing | -0.046179 | 0.002305 | -20.030 | < 2e-16 | < 2e-16 | *** |
| GC content | 0.114300 | 0.003666 | 31.174 | < 2e-16 | < 2e-16 | *** |
| Phased fraction | -0.005726 | 0.005955 | -0.962 | 0.3360 | 0.3840 | |

Supplementary Table 7. The effect of fetal male and female germline expression on all germline mutations, whether or not phased (n=194,356 mutations and 15,220 genes) using a quasipoisson model (dispersion parameter =1.3). Collinearity between male and female germline expression is high (VIF = ~5) but a ridge regression supports coefficient stability.

| | Estimate | Std. Error | t value | Pr(>\|t\|) | FDR q-value |
|---|---|---|---|---|---|
| (Intercept) | -8.634909 | 0.003200 | -2698.639 | < 2e-16 | < 2e-16 *** |
| Fetal male germ. express. | 0.011304 | 0.015114 | 0.748 | 0.45453 | 0.4624 |
| Fetal female germ. express. | 0.039078 | 0.014364 | 2.720 | 0.00653 | 0.00886 ** |
| Male phased fraction | 0.013069 | 0.008447 | 1.547 | 0.12187 | 0.1390 |
| Female phased fraction | 0.025049 | 0.008039 | 3.116 | 0.00184 | 0.00256 ** |
| Replication timing | -0.045952 | 0.002424 | -18.958 | < 2e-16 | < 2e-16 *** |
| GC content | 0.112559 | 0.003634 | 30.977 | < 2e-16 | < 2e-16 *** |

Supplementary Table 8. The effect of GTEx expression on the rate of all phased mutations excluding transitions at CpGs in the male and female germlines using a quasipoisson model (dispersion parameter = 1.36; 15,220 genes). For the intercept and all sex-specific coefficients, the male estimate is shown first (baseline) and then the difference between the male and female coefficients ("female diff.").

| | Estimate | Std. Error | t value | Pr(>\|t\|) | FDR q-value |
|---|---|---|---|---|---|
| (Male Intercept) | -10.303083 | 0.009214 | -1118.239 | < 2e-16 | < 2e-16 *** |
| Male GTEx express. | 0.034742 | 0.018384 | 1.890 | 0.058785 | 0.07129 |
| Intercept (female diff.) | -1.338203 | 0.018836 | -71.044 | < 2e-16 | < 2e-16 *** |
| Male phased fraction | 0.038953 | 0.012383 | 3.146 | 0.001658 | 0.00236 ** |
| Replication timing | -0.098344 | 0.004816 | -20.420 | < 2e-16 | < 2e-16 *** |
| GC content | 0.063428 | 0.007669 | 8.271 | < 2e-16 | < 2e-16 *** |
| GTEx express. (female diff.) | 0.105262 | 0.028166 | 3.737 | 0.000186 | 0.000287 *** |
| GTEx express * phased fraction (male) | -0.007584 | 0.006944 | -1.092 | 0.274782 | 0.2901 |
| Phased fraction (female diff.) | 0.196767 | 0.029069 | 6.769 | 1.32e-11 | 2.59e-11 *** |
| GTEx express.*phased fraction(female diff.) | -0.032816 | 0.016351 | -2.007 | 0.044760 | 0.05547 |

Supplementary Table 9. The effect of pseudobulked fetal germline expression on the rate of all mutations excluding transitions at CpGs in the male and female germlines using a quasipoisson model (dispersion parameter = 1.38; 15,220 genes). For the intercept and all sex-specific coefficients, the male estimate is shown first (baseline) and then the difference between the male and female coefficients ("female diff."). High collineary is present (GVIF = ~4); a ridge regression supports similar coefficients.

| | Estimate | Std. Error | t value | Pr(>\|t\|) | FDR q-value |
|---|---|---|---|---|---|
| (Male Intercept) | -10.307731 | 0.011467 | -898.931 | < 2e-16 | < 2e-16 *** |
| Fetal male germ. express. | 0.020287 | 0.025271 | 0.803 | 0.422 | 0.4375 |
| Female Intercept | -1.246912 | 0.021194 | -58.834 | < 2e-16 | < 2e-16 *** |
| Phased fraction | 0.034030 | 0.024498 | 1.389 | 0.165 | 0.1841 |
| Replication timing | -0.099396 | 0.004894 | -20.310 | < 2e-16 | < 2e-16 *** |
| GC content | 0.060428 | 0.007748 | 7.799 | 6.45e-15 | 1.36e-14 *** |
| Fetal germ. express (female diff.) | 0.258493 | 0.047813 | 5.406 | 6.48e-08 | 1.16e-07 *** |
| Fetal germ. express. * phased fraction (male) | -0.012645 | 0.010911 | -1.159 | 0.246 | 0.2646 |
| Phased fraction (female diff.) | 0.234583 | 0.041021 | 5.719 | 1.08e-08 | 1.99e-08 *** |
| Fetal germ. express.* phased fraction (female diff.) | 0.085449 | 0.020978 | 4.073 | 4.65e-05 | 7.36e-05 *** |

Supplementary Table 10. The effect of GTEx adult gonadal expression on the rate of transitions at CpGs in the male and female germlines using a Poisson model (n=15,220 genes).

| | Estimate | Std. Error | z value | Pr(>\|z\|) | FDR q-value | |
|---|---|---|---|---|---|---|
| (Male Intercept) | -8.139972 | 0.015106 | -538.843 | < 2e-16 | < 2e-16 | *** |
| GTEx express. | -0.057701 | 0.022098 | -2.611 | 0.00902 | 0.01196 | * |
| Intercept (female diff.) | -1.336530 | 0.023204 | -57.600 | < 2e-16 | < 2e-16 | *** |
| Methylation level | 0.364517 | 0.019481 | 18.711 | < 2e-16 | < 2e-16 | *** |
| Phased fraction | 0.043077 | 0.017635 | 2.443 | 0.01458 | 0.01889 | * |
| Replication timing | -0.107972 | 0.008658 | -12.470 | < 2e-16 | < 2e-16 | *** |
| GC content | 0.078534 | 0.012054 | 6.515 | 7.25e-11 | 1.38e-10 | *** |

Supplementary Table 11. The effect of pseudobulked fetal germline expression on the rate of transitions at CpGs in the male and female germlines using a Poisson model (n=15,220 genes).

| | Estimate | Std. Error | z value | Pr(>\|z\|) | FDR q-value | |
|---|---|---|---|---|---|---|
| (Male Intercept) | -8.161716 | 0.015075 | -541.423 | < 2e-16 | < 2e-16 | *** |
| Fetal germ. express. | -0.076614 | 0.032950 | -2.325 | 0.0201 | 0.02546 | * |
| Intercept (female diff.) | -1.323632 | 0.022960 | -57.649 | < 2e-16 | < 2e-16 | *** |
| Phased fraction | 0.037840 | 0.030409 | 1.244 | 0.2134 | 0.2340 | |
| Methylation level | 0.348084 | 0.019537 | 17.817 | < 2e-16 | < 2e-16 | *** |
| Replication timing | -0.101015 | 0.008791 | -11.491 | < 2e-16 | < 2e-16 | *** |
| GC content | 0.052886 | 0.012155 | 4.351 | 1.36e-05 | 2.21e-05 | *** |

Supplementary Table 12. Coefficient estimates from ridge, LASSO and Poisson regression models for the effect of expression levels from adult spermatogonial stem cells and early primary spermatocytes on paternally phased mutations. Poisson GLM coefficients are repeated from Table 1 for ease of comparison.

| | Ridge | LASSO | Poisson GLM |
|---|---|---|---|
| (Intercept) | -10.05153970 | -10.05095754 | -10.05084790 |
| Spermatogonial stem cell | 0.05437827 | 0.05576729 | 0.05831704 |
| Early primary spermatocyte | -0.04791772 | -0.04897051 | -0.05086178 |
| Replication timing | -0.07315945 | -0.07378874 | -0.07399961 |
| GC content | 0.12966689 | 0.13133348 | 0.13173226 |
| Phased fraction | 0.03243393 | 0.03209330 | 0.03227292 |

Supplementary Table 13. The effect of averaged oocyte expression and averaged granulosa cell expression (across five stages of folliculogenesis) from adult females on maternally phased germline mutations using a negative binomial regression model (n=15,220 genes).

| | Estimate | Std. Error | z value | Pr(>\|z\|) | FDR q-value | |
|---|---|---|---|---|---|---|
| (Intercept) | -11.45017 | 0.01349 | -848.616 | < 2e-16 | < 2e-16 | *** |
| Adult oocyte expression | 0.14163 | 0.01904 | 7.437 | 1.03e-13 | 1.58e-13 | *** |
| Adult granulosa cell expression | -0.09841 | 0.01909 | -5.154 | 2.55e-07 | 3.32e-07 | *** |
| Replication timing | -0.11295 | 0.01142 | -9.892 | < 2e-16 | < 2e-16 | *** |
| GC content | 0.17971 | 0.01532 | 11.731 | < 2e-16 | < 2e-16 | *** |
| Phased fraction | 0.04447 | 0.01388 | 3.205 | 0.00135 | 0.00167 | ** |

Supplementary Table 14. The effect of averaged granulosa cell expression (across five stages of folliculogenesis in adult females) on maternally phased germline mutations using a negative binomial regression model (n=15,220 genes).

<table>
<tr><th></th><th>Estimate</th><th>Std. Error</th><th>z value</th><th>Pr(>|z|)</th><th>FDR q-value</th><th></th></tr>
<tr><td>(Intercept)</td><td>-1.144e+01</td><td>1.347e-02</td><td>-849.323</td><td>< 2e-16</td><td>< 2e-16</td><td>***</td></tr>
<tr><td>Adult granulosa cell expression</td><td>-1.779e-02</td><td>1.565e-02</td><td>-1.137</td><td>0.255</td><td>0.265</td><td></td></tr>
<tr><td>Replication timing</td><td>-1.118e-01</td><td>1.147e-02</td><td>-9.749</td><td>< 2e-16</td><td>< 2e-16</td><td>***</td></tr>
<tr><td>GC content</td><td>1.739e-01</td><td>1.532e-02</td><td>11.348</td><td>< 2e-16</td><td>< 2e-16</td><td>***</td></tr>
<tr><td>Phased fraction</td><td>7.653e-04</td><td>1.269e-02</td><td>0.060</td><td>0.952</td><td>0.952</td><td></td></tr>
</table>
